# Seed mycobiome assembly varies across mother trees with contrasting levels of resistance to Dutch elm disease

**DOI:** 10.64898/2026.11.11.765837

**Authors:** Clara Martínez-Arias, Susana Fernández-Quirós, Jesús Rodríguez-Calcerrada, Juan Antonio Martín

## Abstract

Seed-associated fungal communities may influence plant performance from the earliest stages of development, yet their potential association with disease resistance in long-lived forest trees remains poorly understood. Here, we investigated whether *Ulmus minor* trees with different levels of resistance to Dutch elm disease (DED) exhibit consistent differences in their seed-associated mycobiome. We characterized the fungal communities inhabiting samaras from nine *U. minor* genotypes with different levels of DED resistance (i.e. resistant, intermediate and susceptible genotypes) using high-throughput sequencing of the internal transcribed spacer 1 (ITS1) region. We identified 522 fungal amplicon sequence variants (ASVs), including 74 core members of the samara mycobiome. The community was dominated by Dothideomycetes and Tremellomycetes, but both diversity and community composition differed among resistance groups. Susceptible genotypes harbored richer fungal communities, whereas resistant genotypes supported less diverse but more distinct assemblages. Differential abundance analysis identified 21 genera associated with host resistance level, including an enrichment of *Protofenestella* in resistant genotypes and of *Taphrina* in intermediate and susceptible genotypes. Co-occurrence network analysis further revealed contrasting patterns of community organization, with resistant genotypes showing more modular and less connected networks than susceptible genotypes. Together, these findings indicate that DED resistance is associated with distinct patterns of seed mycobiome assembly, diversity and taxonomic composition in *U. minor*. The samara mycobiome may therefore represent a previously overlooked component associated with elm resistance, and provides candidate fungal taxa for future functional validation, with potential applications in elm conservation and mycobiome-informed breeding.

## Introduction

Seeds are vital reservoirs of bacterial and fungal assemblages and a fundamental component of plant–microbe interactions [1]. Seed colonization occurs through three main pathways: (i) an internal maternal route, via the xylem or the non-vascular tissues of the mother plant (maternal vertical transmission); (ii) a floral route, through the stigma (paternal vertical transmission) [2]; and (iii) an external route, from the surrounding environment (horizontal transmission) [3]. Vertically transmitted microbes seem to exhibit evolutionary stability, shaped by natural selection favoring mutualistic interactions [4–6], and may enhance plant survival and fitness under biotic and abiotic stresses [6].

Nevertheless, seed microbiome research has focused mainly on bacteria, while seed-associated fungi remain underexplored, particularly in perennial woody species from natural ecosystems [4]. Plant-associated microbes confer numerous benefits to their hosts [7,8], and fungal endophytes can suppress pathogens directly, through antibiosis, mycoparasitism or niche competition, and indirectly, through nutrient facilitation, hormone modulation or priming of immune responses such as induced systemic resistance [9–12]. Seeds are thus also vectors of microbial inheritance, transmitting fungal endophytes that may influence plant establishment, stress tolerance and disease resistance [13]. The structure and diversity of seed-borne communities depend strongly on host genotype, suggesting that the seed mycobiome may contribute to the differences in disease resistance observed among genotypes [14], and specific taxa have been proposed as indicators of disease-suppressive seed microbiomes [15]. Characterizing the seed mycobiome could therefore provide insights into early-life plant resilience and into the role of the inherited microbiome in long-term resistance against pathogens.

This is particularly relevant for forest species threatened by diseases for which chemical control is unfeasible, such as Dutch elm disease (DED). This vascular wilt disease, caused by *Ophiostoma ulmi* and *O. novo-ulmi*, has devastated *Ulmus* populations in Europe and North America [16,17], where elms were once keystone components of riparian ecosystems [18]. The pathogen, disseminated by elm bark beetles, rapidly colonizes the vascular tissues, causing xylem blockage, foliage wilting and eventually tree death [17]. Identifying and propagating resistant *Ulmus minor* genotypes has therefore become a cornerstone of elm conservation and reforestation. Variation in DED symptoms within natural populations dominated by a single elm genotype, together with differences in resistance among genotypes in controlled trials, suggests that factors beyond host genetics, such as the plant microbiome, may be involved in DED resistance [19,20]. Moreover, DED resistance may be transmitted to a greater extent through the maternal lineage [21–23]; although maternal and paternal effects are difficult to disentangle, transmission of the maternal microbiome to the offspring could be one of the underlying mechanisms. Indeed, certain fungal endophytes isolated from stems of resistant genotypes can mitigate DED by activating the plant immune response, reducing symptoms and buffering the pathogen-induced loss of mycobiome diversity [12,20,24].

Here, we characterize for the first time the samara mycobiome of *U. minor* genotypes with documented differences in susceptibility to *O. novo-ulmi*. We hypothesize that samaras from DED-resistant genotypes harbor fungal assemblages distinct from those of susceptible genotypes, enriched in protective endophytic taxa. Using high-throughput sequencing, we aimed to: (i) compare fungal diversity and community composition in samaras from *U. minor* genotypes with different resistance levels to DED; (ii) identify fungal indicators associated with resistant and susceptible genotypes; and (iii) characterize co-occurrence patterns and potential interactions among fungal taxa. These insights could support microbiome-assisted breeding and sustainable disease mitigation strategies for elm conservation.

## Materials and methods

### Seed sampling and storage

Samaras were collected in April 2020 from nine healthy *Ulmus minor* genotypes in an *ex-situ* conservation plot at the “Puerta de Hierro” Forest Breeding Center (3°45’6” W, 40°27’23” N; 595 m a.s.l., Madrid, Spain). Genotypes were selected to represent a range of DED resistance levels, previously determined in inoculation trials of the Spanish elm breeding program on trees at least 4 years old, using a minimum of 6 ramets per genotype and following the methodology described by [25]. After two consecutive years of artificial inoculation with *O. novo-ulmi*, genotypes VAD2, JCA2 and MDV5 showed crown wilting below 35 % (hereafter resistant genotypes), GRDF3, ABAL1 and MAPD2 between 35 % and 65 % (intermediate genotypes), and MDV1, TOPB1 and MAPD1 above 65 % (susceptible genotypes) **(Table 1)**. All sampled trees grew under very similar micro-site conditions (soil properties, surrounding vegetation and 4 × 4 m spacing) and received drip irrigation in spring and summer to prevent drought stress.

**Table 1:** DED resistance level of the nine *Ulmus minor* genotypes included in the study. Mean foliage wilting percentages were obtained from previous inoculation trials during which an aggressive isolate of *Ophiostoma novo-ulmi* was inoculated in two consecutive years, and refer to 60 days after inoculation. These trials were performed during the regular activities of the Spanish elm breeding program in “Puerta de Hierro” Forest Breeding Center (Madrid, Spain) from 2001 to 2021.

|  |  |  |  | **Susceptibility test** |  |  |
|---|---|---|---|---|---|---|
| **Genotype** | **Provenance in Spain** | **Resistance class** | **Age of sampled tree** | **Foliage wilting after 1st inoculation (%)** | **Foliage wilting after 2nd inoculation (%)** | **N** |
| VAD2 | Valencia, Ademúz | Resistant | 16 | 3.5 ± 2.5 | 8.3 ± 2.7 | 12 |
| JCA2 | Jaén, Cazorla | Resistant | 28 | 24.0 ± 5.4 | 30.8 ± 6.1 | 6 |
| MDV5 | Madrid, Dehesa de Amaniel | Resistant | 22 | 10.7 ± 4.7 | 25.7 ± 4.5 | 12 |
| GRDF3 | Granada, Deifontes | Intermediate | 28 | 23.1 ± 4.6 | 50.9 ± 5.1 | 9 |
| ABAL1 | Albacete, Alatoz | Intermediate | 22 | 43.7 ± 6.4 | 44.2 ± 7.8 | 6 |
| MAPD2 | Málaga, Pedrizas | Intermediate | 22 | 43.5 ± 6.8 | 49.5 ± 6.7 | 10 |
| MDV1 | Madrid, Dehesa de la Villa | Susceptible | 29 | 72.9 ± 4.5 | 94.1 ± 5.5 | 10 |
| TOPB1 | Toledo, Puebla de Montalbán | Susceptible | 29 | 64.5 ± 8.4 | 100 ± 0 | 6 |
| MAPD1 | Málaga, Pedrizas | Susceptible | 28 | 67.5 ± 6.7 | 90.0 ± 7.7 | 8 |

Samaras were collected from one adult tree per genotype (16–29 years old, **Table 1**; 6–12 m in height; 17–32 cm in trunk diameter). Mature samaras at a similar phenological stage (light beige to yellowish-brown pericarp) were sampled with a pole lopper from four fruit-bearing branches per tree, cut from two canopy orientations (northwest and southeast) and from the lower and upper crown. Samples from each tree were pooled, transported to the laboratory at 4 °C, checked manually for fully developed and viable seeds, surface sterilized, frozen in liquid N_2_ and stored at –80 °C. For surface sterilization, 30–35 randomly selected samaras per genotype were shaken for 1 min in 20 mL of filter-sterilized (0.22 µm PES) 1 % Tween 20 solution (Merck KGaA, Darmstadt, Germany), rinsed with autoclaved distilled water, shaken for 1 min in 70 % ethanol, rinsed twice with autoclaved distilled water and air-dried on sterile filter paper in a laminar flow hood.

### DNA extraction and fungal metabarcoding

From each genotype, three subsamples of 10–12 samaras were randomly selected (27 subsamples in total) and ground to a fine powder in a ball mill (Mixer mill MM 400, Retsch GmbH, Haan, Germany). Genomic DNA was extracted from 50 mg of powder with the Norgen Plant/Fungi DNA Isolation Kit (Norgen, Thorold, Canada). After quality control, the first internal transcribed spacer (ITS1) region was amplified with primers ITS1F and ITS2, and libraries were prepared with the Nextera XT Index Kit v2 and sequenced on an Illumina MiSeq platform (2 × 300 bp paired-end reads) by Macrogen Inc. (Seoul, Republic of Korea).

### Analysis of sequencing data and metabarcoding

Raw reads were processed with the DADA2 pipeline v1.22.0 [26] in R (R Development Core Team, Posit Software, PBC, Boston, MA). Primers were removed with CUTADAPT 4.1 [27], chimeras were removed with *removeBimeraDenovo*, and the resulting amplicon sequence variants (ASVs) were taxonomically assigned with the naïve Bayesian classifier [28] against the latest UNITE general fasta release for fungi [29]. ASV counts, taxonomy and sample metadata were combined into a phyloseq object [30], and only fungal sequences were retained. ASVs present in < 5 % of the samples with fewer than five reads were removed as potential artifacts or low-abundance contaminants. For alpha and beta diversity analyses, counts were normalized by scaling with ranked subsampling (SRS) using *srs_p* from the “QsRutils” package [31].

### Data analysis

#### Alpha diversity

Observed richness and the Shannon and Simpson indices were calculated with the “microbiome” package [32]; whereas richness considers only the number of ASVs, the Shannon and Simpson indices also account for their relative abundances [33]. Indices were compared with a nested ANOVA, with genotypes nested within resistance levels, using *aov* with genotype as the error stratum, after checking the normality of residuals (Shapiro–Wilk and Kolmogorov–Smirnov tests, skewness, kurtosis and normal probability plots). We used the functions *shapiro.test* from the R package “stats” to perform the Shapiro-Wilks test and the function *ks.test* from the R package “dgof” for the Kolmogorov-Smirnov test. Skewness and Kurtosis values were obtained with the function *skewness* and *kurtosis* from the R package “moments,” and normal probability plots were drawn using the R packages “ggplot” and “qqplotr.” When the effect of resistance level was significant (P < 0.05), groups were compared with Fisher’s protected least significant difference (LSD) test on genotype means.

#### Beta diversity

Bray–Curtis dissimilarities, calculated on relative abundances, were visualized by principal coordinate analysis (PCoA) with *plot_ordination* (“phyloseq”) and tested by permutational multivariate ANOVA (PERMANOVA) with *adonis2* (“vegan”) *[34]*. Because the three subsamples of each genotype came from a single tree, permutations (9,999) were restricted so that whole genotypes, rather than individual samples, were exchanged among resistance levels (*how* function, “permute” package). Homogeneity of multivariate dispersion was tested with *betadisper* and *permutest* under the same permutation scheme. Pairwise comparisons between resistance levels used the same restricted permutations; as only ten distinct permutations are possible when comparing two groups of three genotypes, pairwise results are reported as effect sizes (R^2^) and interpreted descriptively.

#### Relative abundance of taxa

Relative abundances at the class level were represented for ASVs with a relative abundance higher than 1 % and after filtering free sequence artifacts and low abundant contaminants, using the *tax_glom* and *plot_bar* functions (“phyloseq”).

#### Core and exclusive seed fungal members

Core and exclusive members were identified from relative abundances (*transform_sample_counts*, “phyloseq”) for each resistance level. An ASV was considered part of the core of a group when it exceeded a relative abundance of 0.001 % and a prevalence of 50 % across replicates (*core_members*, “microbiome” [32]). The core shared by all groups and the members exclusive to each group were identified with the *intersect* and *setdiff* functions, respectively, and visualized with Venn diagrams (“ggVennDiagram” [35]).

*Comparison of* U. minor *seed mycobiome with an available* U. minor *stem mycobiome* Seed ASVs were compared with the OTUs previously obtained from stems of adult *U. minor* trees in the same plot [19], a study that included seven of the genotypes analyzed here. Representative ITS sequences of the stem mycobiome were used as BLASTN queries (NCBI BLAST+ v2.17.0+, R-based workflow) against a local database built from the samara sequences. Matches with an E-value ≤ 10⁻⁵, an alignment length ≥ 100 bp and 100 % identity were considered shared taxa and linked to their taxonomic assignment. Because the stem dataset was generated by 454 pyrosequencing, with a lower sequencing depth than Illumina, we assumed that it captured the most abundant and representative members of the stem mycobiome; we therefore restricted the comparison to the taxa shared by both tissues, and their relative abundances are presented for descriptive purposes.

#### Differential abundance

Differences in the relative abundance of fungal genera between resistance levels were tested with the “DESeq2” package [36]. ASVs were agglomerated at the genus level (*tax_glom*, “phyloseq” [30]) and converted with *phyloseq_to_deseq2*, with resistance level as the design factor and subsamples as replicates. As DESeq2 does not accommodate random effects, the non-independence of the three subsamples from each tree could not be explicitly modelled. Pairwise comparisons between resistance levels were performed with Wald tests and P-values were adjusted with the Benjamini–Hochberg method; genera with adjusted P < 0.05 were considered differentially abundant.

#### Co-occurrence networks

Co-occurrence networks were built for each resistance level from pairwise Spearman correlations between ASVs, calculated on the SRS-normalized counts of the nine samples of each level. Only robust and significant correlations (|ρ| > 0.75, P < 0.05) were retained, and positive and negative correlations were interpreted as potential co-occurrence and co-exclusion relationships, respectively. Networks were analyzed with the “igraph” package; modules were identified with the fast-greedy algorithm (checked with the walktrap algorithm), and the modularity of each network was compared with that of 1,000 Erdős– Rényi random networks with the same number of nodes and edges.

## Results

### Fungal community and taxonomic composition

Sequencing of the 27 subsamples yielded 4,233,864 reads (mean 156,809 per subsample; range 146,543–193,866), distributed in 1,316 ASVs, of which 522 were retained after filtering. These were assigned to 253 species, 166 genera, 101 families, 51 orders and two phyla, Ascomycota (70 % of reads) and Basidiomycota (30 %). Dothideomycetes, Tremellomycetes, Taphrinomycetes and Microbotryomycetes dominated in all genotypes (**Fig. 1**). Although the same major classes were detected at all resistance levels, their relative abundances differed markedly among them.

**Fig. 1.**
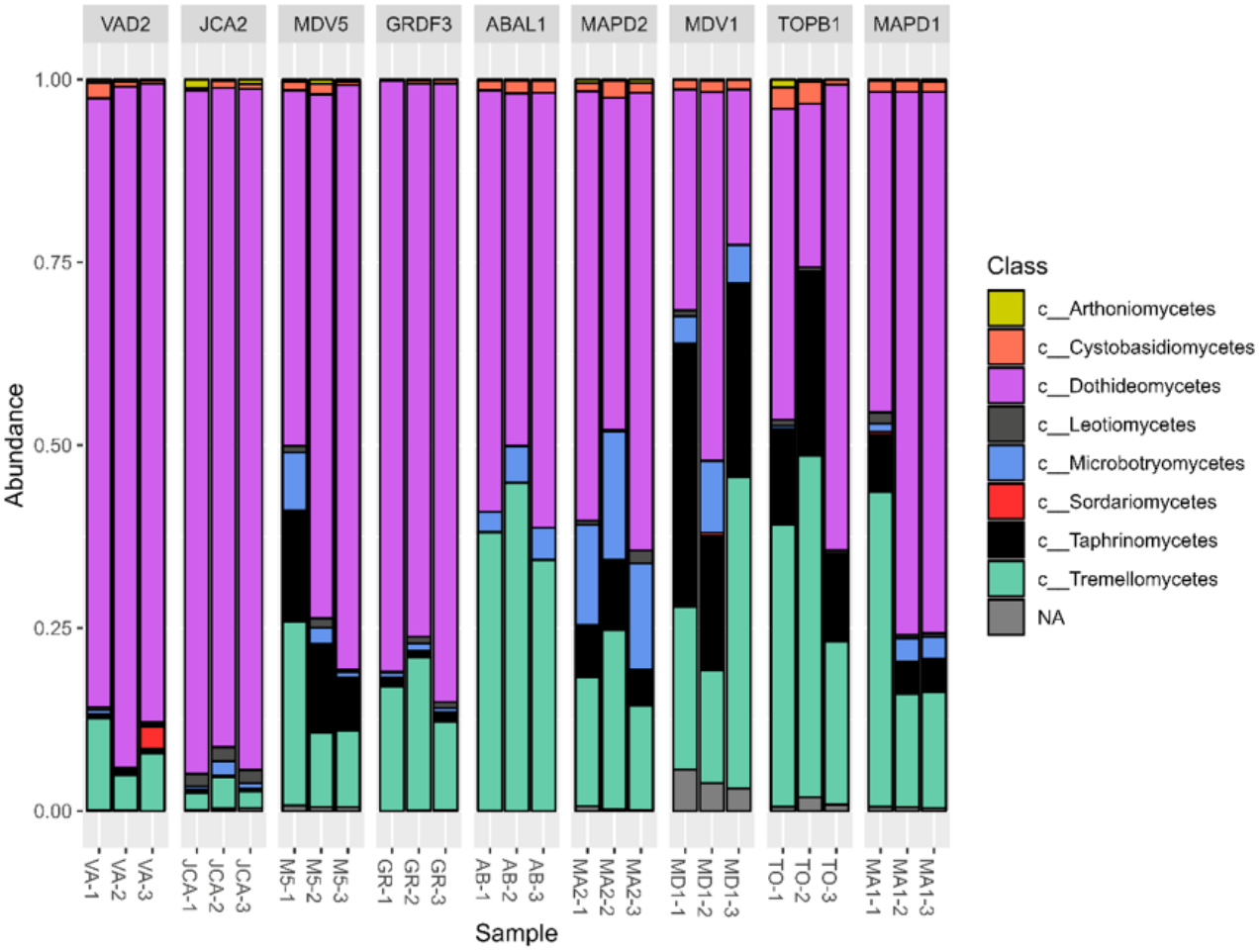
Class-level relative abundance of the samara fungal communities in the three replicates of samaras from nine *Ulmus minor* genotypes with contrasting levels of resistance to Dutch elm disease (DED). VAD2, JCA2 and MDV5 are DED-resistant genotypes; GRDF3, ABAL1 and MAPD2 are DED-intermediate genotypes; and MDV1, TOPB1 and MAPD1 are DED-susceptible genotypes.

### Biodiversity analysis and shared seed fungal members between elm resistance levels

Alpha diversity differed among resistance levels, particularly for observed richness and Shannon diversity (**Fig. 2A**). Richness differed significantly among all three levels, being highest in susceptible genotypes, intermediate in resistant genotypes and lowest in intermediate genotypes. Shannon diversity was significantly higher in susceptible genotypes than in resistant and intermediate ones, which did not differ from each other. The Simpson index also tended to be higher in susceptible genotypes, although differences were marginally non-significant (P = 0.07; **Table 2**). Overall, susceptible genotypes supported richer and more diverse fungal communities than resistant and intermediate genotypes.

**Fig. 2.**
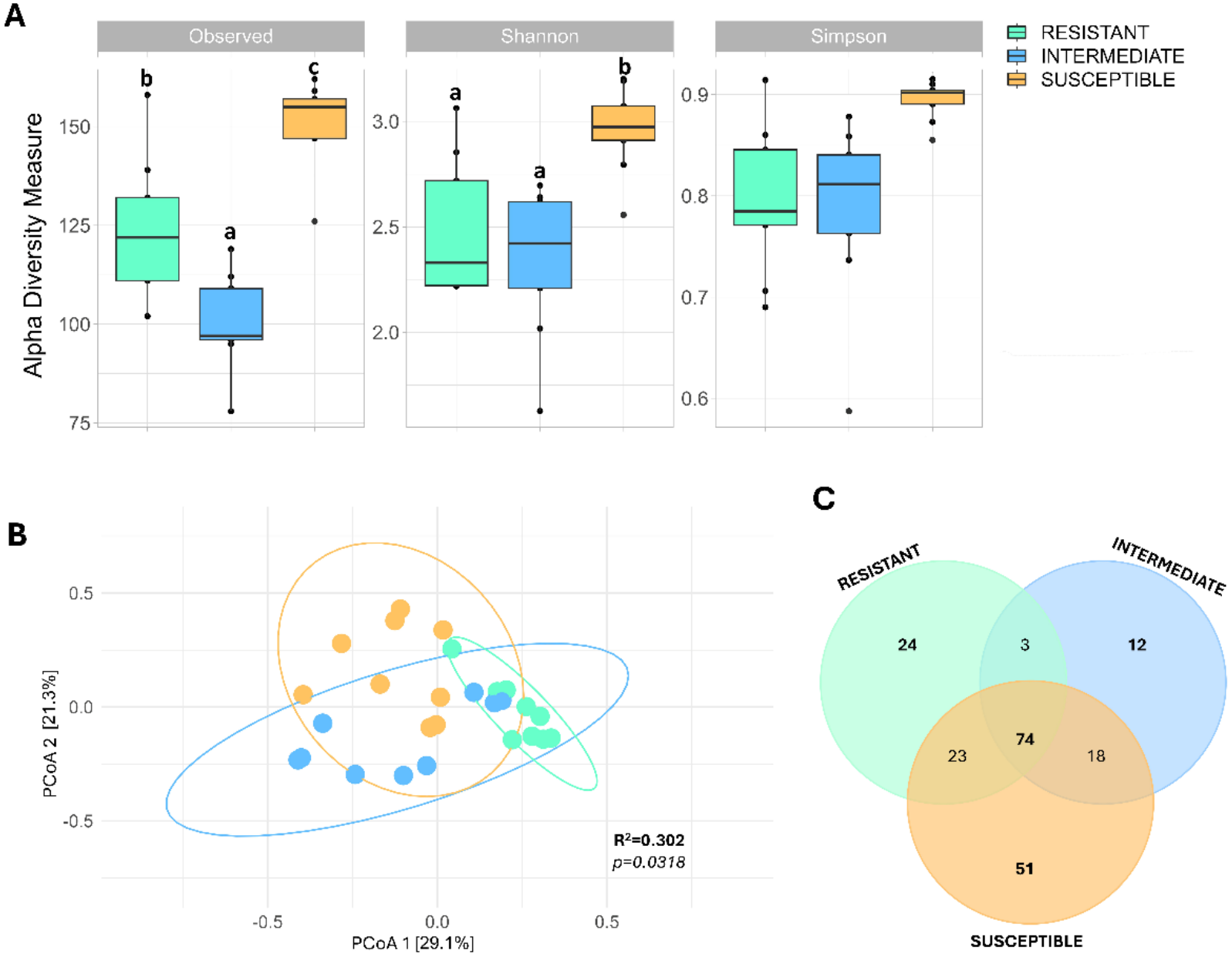
**(A)** Alpha diversity of the *Ulmus minor* samara mycobiome based on observed amplicon sequence variant (ASV) richness, Shannon diversity and Simpson diversity indices according to DED-resistance level. Different letters indicate significant differences between groups according to Fisher’s post hoc test (*P* < 0.05) after nested one-way ANOVA analysis in which resistance level was tested against the variation among genotypes. **(B).** Differences in fungal community composition of the *Ulmus minor* samara mycobiome according to DED-resistance level. Principal coordinate analysis (PCoA) of mycobiome composition based on Bray-Curtis dissimilarities among the three DED-resistance levels. *R*² and *P* values indicate the results of the PERMANOVA analysis. **(C)** Venn diagram showing shared and exclusive amplicon sequence variants (ASVs) among the three DED-resistance levels.

**Table 2:**
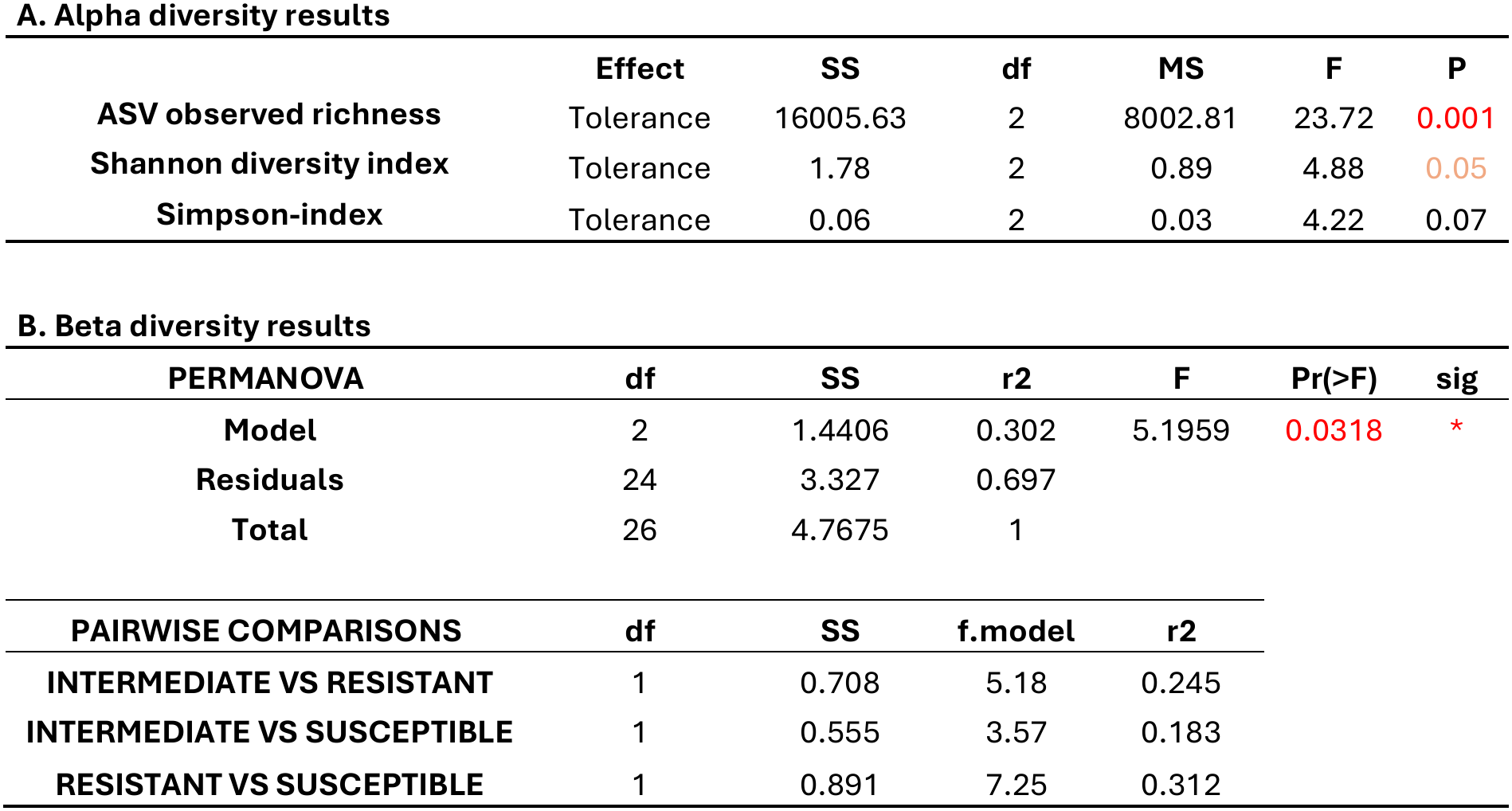
Statistical analyses of alpha and beta diversity of the *Ulmus minor* samara mycobiome. (A) Alpha diversity, measured as observed amplicon sequence variant (ASV) richness and Shannon and Simpson indices. The effect of Dutch elm disease (DED) resistance level was assessed with nested ANOVAs in which resistance level was tested against the variation among genotypes within resistance levels. (B) Beta diversity, analyzed with permutational multivariate analysis of variance (PERMANOVA) on Bray–Curtis dissimilarities, with 9,999 permutations of whole genotypes among resistance levels. Pairwise comparisons are reported as effect sizes (R²). SS, sum of squares; df, degrees of freedom; MS, mean square; F, F statistic; R², proportion of variance explained. Red letters indicate significant effects (P < 0.05).

| **A. Alpha diversity results** |  |  |  |  |  |  |
|---|---|---|---|---|---|---|
|  | **Effect** | **SS** | **df** | **MS** | **F** | **P** |
| **ASV observed richness** | Tolerance | 16005.63 | 2 | 8002.81 | 23.72 | 0.001 |
| **Shannon diversity index** | Tolerance | 1.78 | 2 | 0.89 | 4.88 | 0.05 |
| **Simpson-index** | Tolerance | 0.06 | 2 | 0.03 | 4.22 | 0.07 |
| **B. Beta diversity results** |  |  |  |  |  |  |
| **PERMANOVA** | **df** | **SS** | **r2** | **F** | **Pr(>F)** | **sig** |
| **Model** | 2 | 1.4406 | 0.302 | 5.1959 | 0.0318 | * |
| **Residuals** | 24 | 3.327 | 0.697 |  |  |  |
| **Total** | 26 | 4.7675 | 1 |  |  |  |
| **PAIRWISE COMPARISONS** | **df** | **SS** | **f.model** | **r2** |  |  |
| **INTERMEDIATE VS RESISTANT** | 1 | 0.708 | 5.18 | 0.245 |  |  |
| **INTERMEDIATE VS SUSCEPTIBLE** | 1 | 0.555 | 3.57 | 0.183 |  |  |
| **RESISTANT VS SUSCEPTIBLE** | 1 | 0.891 | 7.25 | 0.312 |  |  |

PERMANOVA revealed significant differences in community composition among resistance levels, which explained 30.2 % of the variation (R^2^ = 0.30; P = 0.0318; **Fig. 2B**, **Table 2**). Multivariate dispersion did not differ among groups (P = 0.37), indicating that these differences reflect shifts in composition rather than in within-group variability. Pairwise effect sizes were largest between resistant and susceptible genotypes (R^2^ = 0.31), moderate between resistant and intermediate genotypes (R^2^ = 0.25) and smallest between intermediate and susceptible genotypes (R^2^ = 0.18; **Table 2**). Consistently, resistant genotypes clustered together along the first PCoA axis (29.1 % of the variation), susceptible genotypes were displaced towards the opposite end, and intermediate genotypes spanned a broader range of the ordination space (**Fig. 2B**).

Of the 205 ASVs retained in the core analysis, 74 (36 %) formed the core seed mycobiome (**Fig. 2C; Supp. Table 1**), which was dominated by Tremellomycetes (40.5 %) and Dothideomycetes (35.5 %). Susceptible genotypes had the highest number of exclusive ASVs (51), followed by resistant genotypes (24).

### Comparison with the stem mycobiome of trees from the same plot

Forty-five ASVs/OTUs were shared between the samara and stem mycobiomes (**Fig. 3; Supp. Table 2**). In both tissues, these shared taxa were dominated by Dothideomycetes, although Lecanoromycetes were substantially less abundant and Tremellomycetes more abundant in samaras (**Fig. 3A**). At the genus level, *Aureobasidium* was a major constituent in both tissues, while *Protofenestella*, *Cladosporium* and *Alternaria* were well represented in samaras (**Fig. 3B**). The yeasts *Buckleyzyma* (Cystobasidiomycetes) and *Genolevuria* (Tremellomycetes) were also notably present in both tissues.

**Fig. 3.**
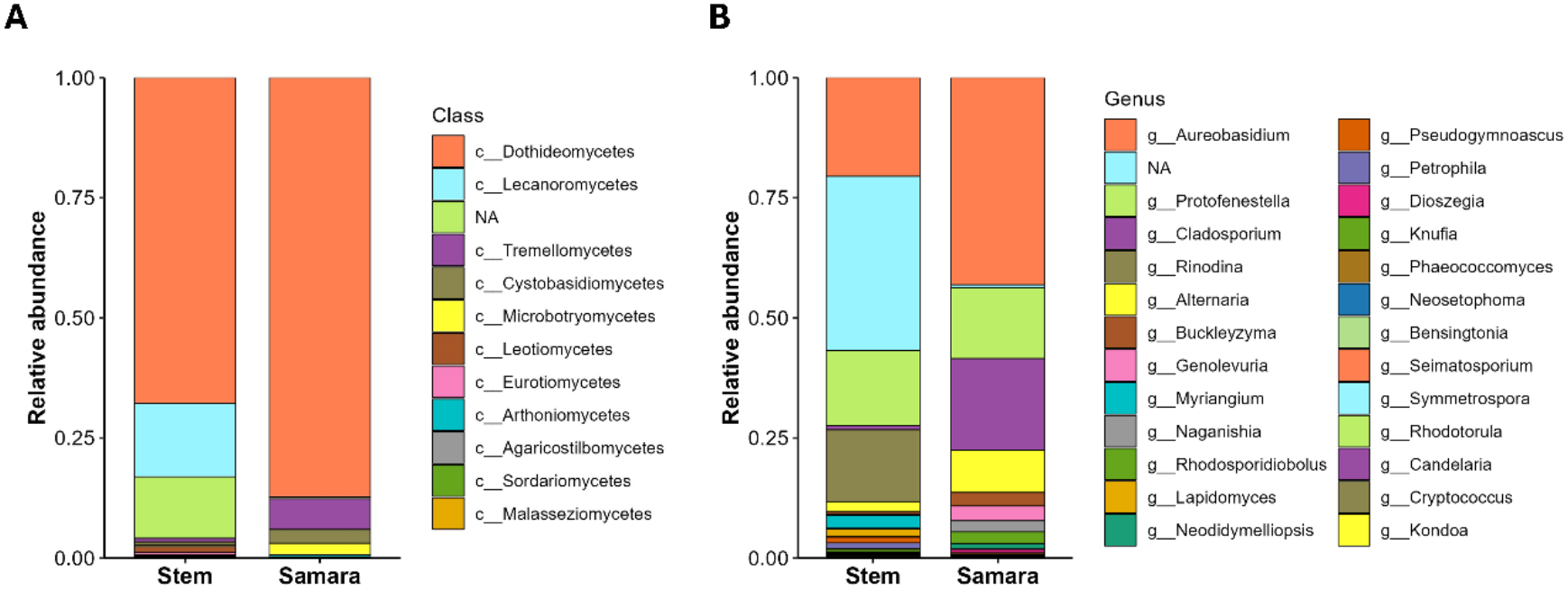
Class- and genus-level relative abundance of the 45 amplicon sequence variants (ASVs) shared between all samples of the *Ulmus minor* samara mycobiome analyzed in this study and the stem mycobiome reported by Macaya-Sanz et al. (2023) [19].

### Differentially abundant fungi between elm resistance levels

DESeq2 identified 21 genera as differentially abundant in at least one of the three pairwise comparisons between resistance levels (**Fig. 4; Supp. Table 3**). The most abundant of these, with similar abundances across the three genotypes within each resistance level, were *Protofenestella*, *Udeniomyces*, *Taphrina*, *Aureobasidium*, *Filobasidium*, *Cryptococcus*, *Genolevuria*, *Buckleyzyma* and *Kwoniella* (**Fig. 4**). The potential plant pathogen *Taphrina* and the yeasts *Filobasidium* and *Cryptococcus* were significantly enriched in intermediate and susceptible genotypes relative to resistant ones, whereas *Aureobasidium*, *Buckleyzyma*, *Kwoniella* and *Genolevuria* were enriched only in susceptible genotypes. Conversely, *Protofenestella* was significantly more abundant in resistant than in intermediate and susceptible genotypes. Thus, DED resistance was associated with a distinct seed mycobiome signature, including a reduction of potentially pathogenic and opportunistic taxa such as *Taphrina*.

**Fig. 4.**
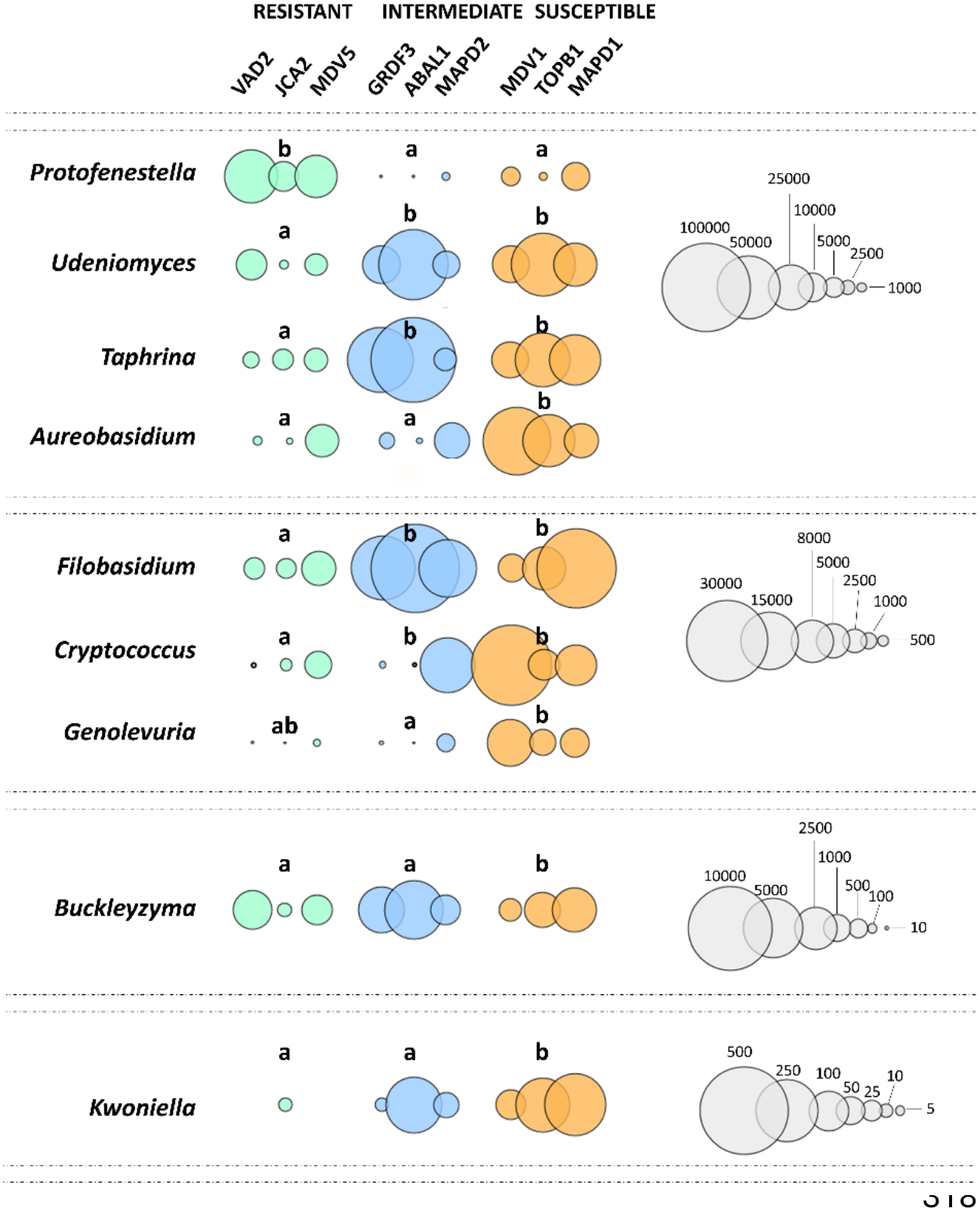
Differentially abundant fungal genera among the samaras of the three DED- resistance levels (resistant, intermediate and susceptible) identified with DESeq2. Different letters indicate significant differences between resistance levels based on pairwise Wald tests (Benjamini–Hochberg adjusted P < 0.05). Bubbles represent the mean number of DESeq2-normalised reads of each genus in each genotype, colored by resistance level. Owing to the large differences in read counts among genera, genera are grouped into blocks, each with its own bubble size scale (right).

### Co-occurrence network analysis of the elm seed mycobiome

Exploratory co-occurrence networks differed in structure among resistance levels (**Fig. 5**; **Table 3**). The susceptible network was the most densely connected (89 edges among 34 ASVs; density = 0.16), whereas the resistant network was the least connected (68 edges among 35 ASVs) and was split into three components; the intermediate network showed intermediate values (92 edges among 36 ASVs). Negative associations were present in all networks and accounted for 29.4 %, 47.8 % and 34.8 % of the edges in resistant, intermediate and susceptible genotypes, respectively. The networks of resistant and intermediate genotypes were more modular than expected by chance (modularity = 0.44 and 0.36, compared with 0.38 and 0.32 in random networks; P = 0.019 and 0.029, respectively), whereas that of susceptible genotypes was not (0.29 compared with 0.31; P = 0.78).

**Fig. 5.**
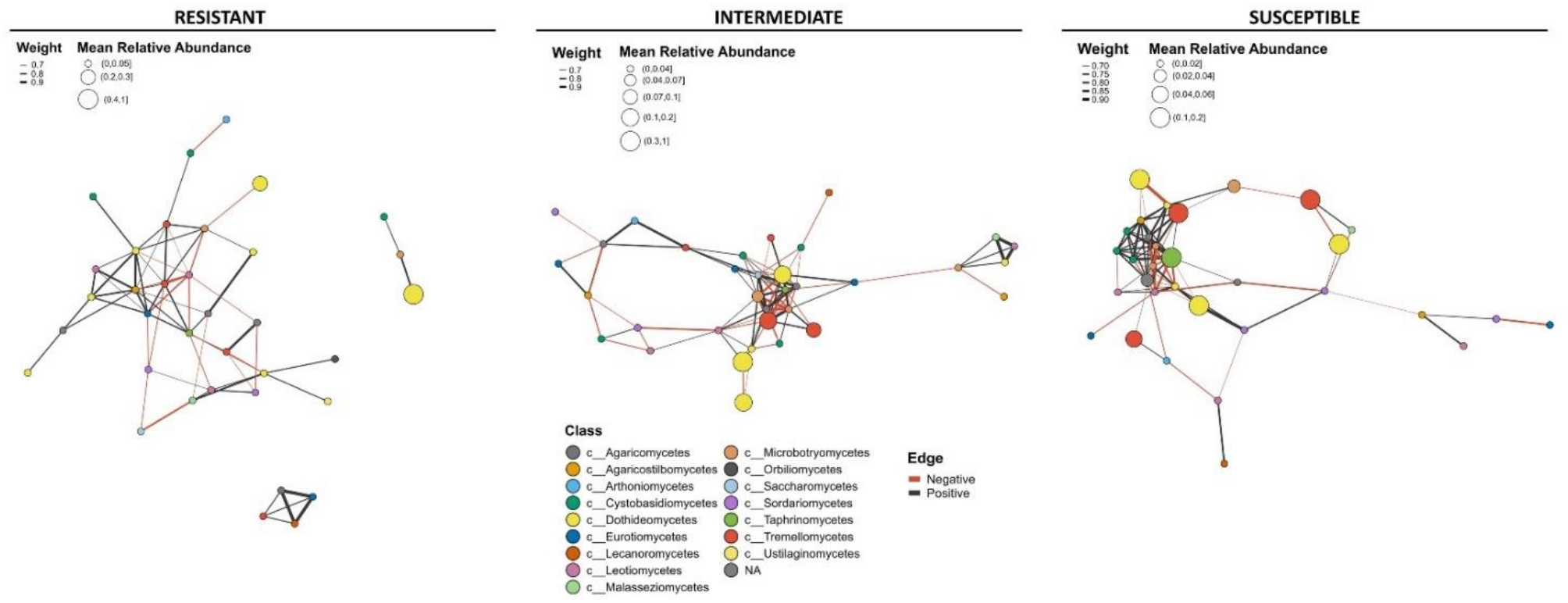
Spearman co-occurrence networks of the samara mycobiome of *Ulmus minor* genotypes with contrasting levels of DED resistance. Networks are shown separately for resistant, intermediate and susceptible genotypes. Each node represents a fungal taxon, with node size proportional to its mean relative abundance and node color indicating fungal class. Edges represent significant associations between fungal taxa, with positive and negative correlations shown in black and red, respectively. Only strong associations (|*ρ*| > 0.75) are shown.

**Table 3:**
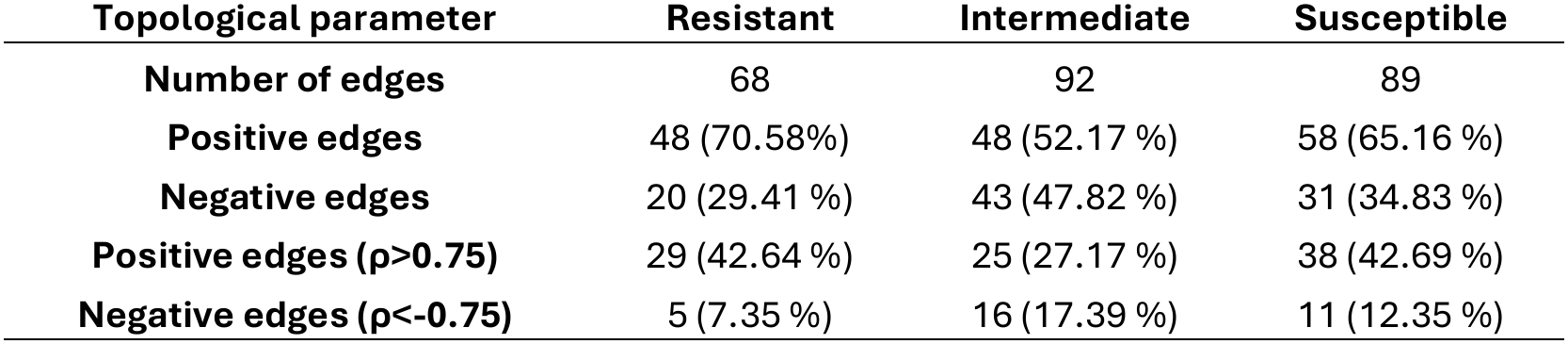
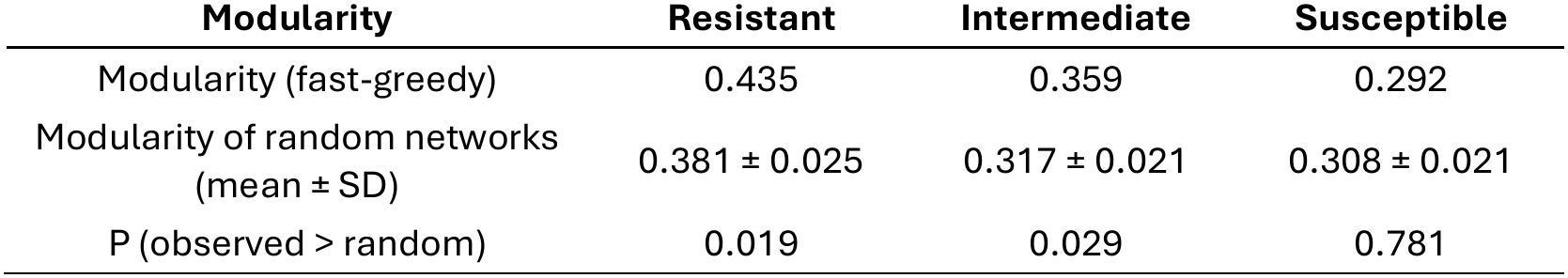
Topological parameters of Spearman co-occurrence networks constructed from the fungal members of the *Ulmus minor* samara mycobiome according to host DED resistance. Networks were constructed separately for resistant, intermediate and susceptible genotypes. Number of edges represents the total number of significant associations between fungal members. Positive and negative edges indicate positive and negative Spearman correlations, respectively, based on an absolute correlation coefficient threshold of |*ρ*| > 0.75 and *P* < 0.05. Modularity of the co-occurrence networks of each resistance level compared with that of 1,000 Erdős–Rényi random networks with the same number of nodes and edges. P indicates the proportion of random networks with a modularity equal to or higher than the observed value.

## Discussion

This study provides the first characterization of the samara mycobiome of *U. minor* genotypes with contrasting resistance to DED. By analyzing intact samaras, including both the seed and the pericarp, we also captured fungi from the pericarp, a maternal tissue that can act as a microbial reservoir and transfer microbes to the emerging seedling during germination. Mycobiome composition, diversity and co-occurrence patterns varied among samaras from mother trees with different resistance levels. As all mother trees shared similar environmental conditions, these differences reinforce the view that the seed microbiome can be strongly shaped by host genotype.

### The samara mycobiome shares core fungal taxa with the stem endosphere

The samara mycobiome was dominated by Dothideomycetes, Tremellomycetes, Taphrinomycetes and Microbotryomycetes, with Arthoniomycetes, Cystobasidiomycetes, Leotiomycetes and Sordariomycetes at lower abundances (**Fig. 1**). Its core included *Alternaria*, *Aureobasidium* and *Cladosporium* (Dothideomycetes) and basidiomycetous yeasts such as *Naganishia*, *Vishniacozyma*, *Dioszegia* and *Filobasidium* (Tremellomycetes) (**Fig. 2C, Supp. Table 1**). *Alternaria*, *Cladosporium* and yeasts such as *Vishniacozyma* are consistently identified as ubiquitous members of the global seed core mycobiome across plant species [37,38].

The comparison with the stem mycobiome of *U. minor* trees from the same plot [19] revealed shared taxa able to colonize both niches. The overlap was most pronounced at higher taxonomic ranks, with Dothideomycetes, Tremellomycetes and Cystobasidiomycetes dominating across tissues. The 45 shared ASVs/OTUs were dominated by *Aureobasidium*, *Protofenestella*, *Cladosporium*, *Alternaria*, *Buckleyzyma* and *Genolevuria*, and included *Knufia* (Eurotiomycetes) and *Rhodotorula* (Sporidiobolales), previously isolated, cultured and functionally characterized in *U. minor* [12,24,39].

Although our design cannot discriminate between vertically and horizontally transmitted taxa, this overlap suggests that a substantial proportion of stem endophytes can also colonize samaras, increasing their likelihood of becoming primary colonizers of emerging seedlings [40]. The recurrence of these taxa across independent studies and tissues suggests that certain members of the *U. minor* mycobiome are systematically maintained, and the prevalence of yeasts and yeast-like fungi agrees with their frequent presence in seed tissues [41,42]. While direct experimental validation (e.g. tracking fluorescently labeled strains) is needed to confirm seed-borne inheritance, our findings provide candidate seed-transmitted fungi.

### DED resistant elms are associated with reduced seed mycobiome diversity

Alpha and beta diversity differed between DED-resistant and susceptible genotypes (**Fig. 2A, B**), suggesting that the host genetic background exerts a selective pressure on seed fungal community assembly. Genotype-dependent seed mycobiomes have also been described in *Triticum* spp., *Brassica napus*, *Cucurbita pepo* and *Quercus petraea* [43–46]. In *U. minor*, distinct stem fungal communities have likewise been associated with different levels of DED resistance, indicating that genotype-dependent filtering may operate throughout plant development [19,47].

Intriguingly, susceptible genotypes showed higher fungal richness and diversity than resistant ones (**Fig. 2A**). While this contrasts with the classic ecological paradigm associating higher diversity with greater community stability [48], it matches the findings by Martín et al. (2013) [47] with culturable *U. minor* xylem endophytes and observations in other pathosystems. For instance, seeds of *Brassica napus* cultivars resistant to *Plasmodiophora brassicae* harbor less diverse communities than susceptible ones [45], and cultivated *Triticum aestivum*, which tends to be more susceptible to diseases than its wild relatives, harbors more diverse and variable fungal assemblages [46]. We hypothesize that resistant *U. minor* genotypes do not rely on a broad, unselective microbial consortium, but selectively recruit a more specialized community enriched in key taxa that confer resilience against biotic stress [49]. These early colonizers could occupy key ecological niches and create a disease-suppressive seed microbiome acting as a primary line of defense from the onset of germination [15]. Conversely, the higher diversity in seeds of susceptible genotypes points to a more permissive seed environment and a lower capacity of the host to discriminate among beneficial, neutral and potentially harmful fungi [50].

Host defense activation can act as an active filter of microbiome assembly, with immune signaling and specialized metabolites shaping which microbes establish and persist [51]. In *U. minor*, susceptible genotypes show weak or delayed defense activation against *O. novo-ulmi* [52], a permissive response associated with more frequent shifts in the stem mycobiome [20]. This effect may extend to seeds through the maternal mycobiome or through defense mechanisms operating within seeds. Moreover, maintaining a more diverse fungal community may entail additional costs for the host, potentially compromising the resilience of the new plant [53].

### DED-resistant genotypes selectively filter fungi colonizing samaras

The fungi exclusive to each resistance level (**Fig. 2C, Supp. Table 1**) further support a potential contribution of the seed mycobiome to disease resistance in seedlings. The exclusive mycobiome of resistant genotypes included putative beneficial taxa, but also taxa considered pathogens in other systems, such as *Fusarium commune* and species of *Stemphylium* and Didymellaceae [54–56], none of which has been documented as a pathogen of *Ulmus*.

Among putative beneficial taxa exclusive to resistant genotypes were the basidiomycetous yeasts *Vishniacozyma dimennae* and *Sporobolomyces reniformis*, well-documented epiphytic and endophytic colonizers of the phyllosphere [41,57] and core members of the global seed mycobiome [37,38]. *Vishniacozyma* in seeds has been correlated with high levels of disease suppression against *Globisporangium ultimum* in spinach and against *Tilletia* and *Fusarium* in wheat [15,41], and it is more abundant in *Fraxinus excelsior* genotypes less susceptible to ash dieback, where it may participate in pathogen antagonism [58]. The exclusive presence of *Cladosporium cladosporioides*, a vertically transmitted endophyte that may enhance seed vigor [59], further supports the idea of a protective consortium that could limit pathogen establishment through niche preemption, nutrient competition and mycoparasitism [41,60].

This putative consortium also included the dark septate endophytes *Pseudogymnoascus appendiculatus* and *Knufia cryptophialidica*, which typically enhance tolerance to abiotic and oxidative stress, critical during germination and early establishment [61–63]. The detection of *Knufia* and *Rhodotorula*, previously isolated from stems of resistant elms [19], reinforces the hypothesis of transmission of protective endosymbionts.

In contrast, the exclusive presence of phytopathogens such as *Taphrina carpini*, *Botrytis* sp. and *Didymella glomerata* in susceptible genotypes points to a more permissive seed environment. Although only *Taphrina* species are known pathogens of *Ulmus*, *Botrytis* and *Didymella* species are key indicators of low-performing seed lots that fail to suppress damping-off [15]. Together with the higher number of exclusive taxa, this suggests that susceptible genotypes exert weaker filtering against opportunistic fungi, resulting in a broader seed fungal community with a greater presence of potential opportunistic parasites [64].

### Differential abundance of taxa across the resistance gradient

Differential abundance analysis further supports host-mediated filtering of the seed mycobiome (**Fig. 4**). Resistant genotypes were enriched in *Protofenestella*, a genus within the Dothideomycetes, a class frequently dominant in seed-associated communities and repeatedly implicated in biocontrol and plant protection [6]. Dothideomycetous endophytes have been associated with DED resistance in adult elm tissues [19], and their beneficial roles have been validated *in vivo [12,24,3S]*. Their detection in samaras may indicate transmission to juvenile plants, whose early microbiome is largely seed-derived [65]. In contrast, seeds of susceptible and intermediate genotypes were enriched in *Taphrina*, a genus associated with leaf galls and deformities in woody hosts [66], again pointing to a less restrictive seed environment open to colonization by opportunists and latent pathogens [46].

The enrichment of *Aureobasidium*, *Filobasidium*, *Cryptococcus*, *Genolevuria*, *Buckleyzyma* and *Kwoniella* in susceptible genotypes also reflects the accumulation of a wider range of taxa than in the more restricted assemblages of resistant trees. However, *Filobasidium* and *Aureobasidium* are ubiquitous core members of seeds with documented biocontrol properties based on nutrient competition and secondary metabolite production [38,67], and species of *Aureobasidium*, *Filobasidium* and *Genolevuria* belong to the core stem mycobiome of *U. minor* [19]. Thus, although seeds of susceptible trees also harbor potentially beneficial taxa, their overrepresentation suggests weaker directional filtering by the host.

### Mycobiome network architecture differs across the DED resistance gradient

Co-occurrence networks revealed fundamental differences in mycobiome architecture across the resistance gradient (**Fig. 5**). The dense, highly interconnected and poorly modular network of susceptible genotypes is consistent with a weak subdivision into ecologically cohesive subcommunities [68,69], which may render the mycobiome vulnerable to cascading disruptions following pathogen invasion, as perturbations propagate rapidly through such communities [70,71]. Resistant genotypes displayed a more modular architecture, with a cohesive core cluster and topologically isolated subgroups of taxa, as also observed in coffee varieties resistant to coffee leaf rust [72].

Modularity may reflect the organization of the community into stable consortia and niche differentiation that reduces competition and promotes complementary functions [68], potentially enhancing community resilience and the ability of the host to resist pathogen establishment, as described for disease-suppressive soils [73].

Together with the diversity results, these network topologies suggest divergent ecological strategies: susceptible genotypes harbor diverse but weakly structured and permissive fungal assemblages, potentially reflecting reduced host-mediated filtering, whereas resistant genotypes host more constrained and modular assemblages, suggesting stronger host selection of a curated subset of taxa.

### Conclusions

The distinct samara mycobiomes of *U. minor* genotypes suggest that the seed mycobiome may contribute to the assembly of the plant mycobiome, with potential physiological consequences throughout the tree’s lifespan. Our findings support the hypothesis that DED resistance is linked to the transmission or acquisition of a curated, highly structured and potentially protective seed mycobiome, which could shape seedling mycobiome assembly through priority effects, pre-empting niches before *O. novo-ulmi* colonizes the vascular system. Functional validation through synthetic communities and controlled inoculation trials is still needed to establish causality. Nevertheless, the genotype-specific fungal indicators and co-occurrence patterns identified here open promising avenues for mycobiome-assisted breeding and sustainable elm conservation.

## Acknowledgements

We thank the Spanish elm breeding program and Puerta de Hierro Forest Breeding Centre (Ministerio para la Transición Ecológica y el Reto Demográfico) for providing the plant material and Jorge Domínguez for technical assistance.

This study was funded by the project AGL-2015-66952-R (MINECO/FEDER). C.M.-A. was supported by a ‘FPI’ pre-doctoral fellowship from the Spanish Ministry of Economy and Competitiveness during 2017-2021.

## Author Contributions

C.M.-A. performed the experiments and laboratory analysis, analyzed and interpreted the data, and wrote the main manuscript text. S.F.-Q. contributed to laboratory and field experiments, data collection, and data interpretation. J. R.-C. and J.A.M. conceived and designed the study, contributed to data interpretation, and manuscript writing and revision. All authors reviewed, edited, and approved the final manuscript.

## Data Availability

Raw sequences of the mycobiome are available online in NCBI under the BioProject code PRJNA1548562.

